# The probiotic effectiveness in experimental colitis is correlated with gut microbiome and host genetic features

**DOI:** 10.1101/340331

**Authors:** Sharmila Suwal, Qiong Wu, Wenli Liu, Qingya Liu, Hongxiang Sun, Ming Liang, Jing Gao, Bo Zhang, Yanbo Kou, Zhuanzhuan Liu, Yanxia Wei, Yugang Wang, Kuiyang Zheng

**Author notes:** **Correspondence:** Yugang Wang, or Kuiyang Zheng, Xuzhou Medical University, Tongshan Road #209, Xuzhou, Jiangsu Province, P.R. China, 221004.

## Abstract

Current evidence to support extensive use of probiotics in inflammatory bowel disease is limited and factors contribute to the inconsistent effectiveness of clinical probiotic therapy are not completely known. Here, as a proof-of-concept, we utilized *Bifidobacterium longum* JDM 301, a widely used commercial probiotic strain in China, to study potential factors that may influence the beneficial effect of probiotics in experimental colitis. We found that the probiotic therapeutic effect was varied across individual mouse even with the same genetic background and consuming the same type of food. The different probiotic efficacy was highly correlated with different microbiome features in each mouse. Consumption of a diet rich in fat can change the host sensitivity to mucosal injury-induced colitis but did not change the host responsiveness to probiotic therapy. Finally, the host genetic factor TLR2 was required for a therapeutic effect of *B. longum* JDM 301. Together, our results suggest that personalized microbiome and genetic features may modify the probiotic therapeutic effect.

## Introduction

It has long been recognized that the microbiota in the gut can impact many aspects of the host biology. Live microbes that confer health benefits to the host are often called probiotics. Consumption of probiotics in various forms like yogurt or other fermented dairy products, as dietary supplements and other functional foods, has become more and more popular. Probiotics are also claimed to have therapeutic benefits across a broad range of disorders including diseases in the gastrointestinal tract (1–3). Inflammatory bowel disease (IBD), including Crohn’s disease (CD) and ulcerative colitis (UC), is a multi-factorial complex intestinal disorder with the highest prevalence in western countries (4–6). Probiotics are recommended by physicians as adjunctive therapy to treat IBD (7). Despite their popularity, the current evidence to support extensive use of probiotics in IBD is limited. Results from clinical trials are mixed, with some studies showing an improvement in maintenance of remission or induction of remission with probiotics while other trials have failed to show any benefit effect (8, 9). The reason behind the various outcomes of probiotic effectiveness in treating IBD is not clear.

It is now widely acknowledged that the gut microbiome together with the host genetic factors significantly contribute to the pathogenesis of IBD (10). Gut microbiota plays significant roles by preventing pathogen colonization (11), shaping the immune system (12, 13), stimulating the production of gastrointestinal hormones (14), regulating brain behavior through the production of neuroactive substances and fermentation of non-digestible carbohydrates producing short chain fatty acids (SCFAs) (15, 16). Most recently, the microbiome is also emerging as contributing factor to interindividual variability in all aspects of a disease (17). However, whether the gut microbiota contributes to the person-to-person differences in response to probiotic therapy remains largely unknown.

*Bifidobacterium longum* JDM 301, isolated from healthy infants, is a widely used commercial probiotic strain in China (18). Our previous study demonstrated that *B. longum* JDM 301 can prevent *Clostridium difficile* infection (CDI) in mice (19). In the present study, we used *B. longum* JDM 301 as a proof-of-concept to test factors that could potentially influence the therapeutic effect of probiotics in experimental colitis. Our data demonstrate associations of the gut microbiome and host genetic factor to interindividual variability in probiotic biotherapeutic responses. Our results suggest personalized strategies are needed for the success of probiotics therapies.

## Materials and Methods

### Animals

5 to 6-week-old male C57Bl/6J mice were obtained from Shanghai Laboratory Animal Research Center, Shanghai or Beijing Vital River Laboratory Animal Technology Co., Ltd., Beijing. *TLR2^-/-^* were originally purchased from Model Animal Research Center of Nanjing University and maintained under specific pathogen-free (SPF) conditions with a 12 hours light and 12 hours dark cycle and had free access to diet and drinking water. All animal procedures were done following the institutional guidelines and approved by the Animal Care and Use Committee of the University.

### Diet

WT and *TLR2^-/-^* mice were fed with standard chow diet for the first week of arrival to the laboratory. Then the mice were randomly divided into two groups. One group of mice was fed with standard chow diet (ND) and the other group of mice was fed with high-fat diet (HFD), 60% energy from fat (Ke Ao Xie Li Co. Ltd., Beijing). HFD feeding was continued for a total of 6 weeks.

### Colitis induction

Colitis was induced using DSS (Molecular Weight = 36,000~50,000) (MP Biomedicals, Santa Ana, CA, USA). For mice under ND group (5-8 mice), DSS treatment was started when the mice became 8 weeks of age. For mice under HFD group (5-8 mice), DSS induced colitis was started after 6 weeks of HFD feeding. Both groups were given 3% DSS (w/v) in drinking water for 7 days followed by 3 days of recovery period during which sterilized drinking water was supplied. The control group (3-5 mice) fed with either ND or HFD received only sterilized drinking water throughout the experiment.

### Preparation of *B. longum* JDM 301 for inoculation to mice

The *B. longum* JDM 301 was originally isolated from a commercial probiotic product from China (18). The frozen glycerol stock of *B. longum* JDM 301 was thawed and then plated on MRS agar plate. The plate was incubated anaerobically overnight. The next day, a single bacterial colony was inoculated in a tube containing 3-5 ml of MRS broth and was incubated for 16-24 hours anaerobically. The total number of bacteria for each mouse used each time was 1×10^9^ colony forming unit (cfu).

### Colonic Tissue Collection and Processing

The colons tissues were collected and fixed in 4% (w/v) paraformaldehyde (pH 7.0), dehydrated by increasing concentrations of ethanol, and embedded in paraffin for histological studies.

### Histological Examination

For histological grading three different parameters were considered, severity of inflammation (based on polymorphonuclear neutrophil infiltration; 0–3: none, slight, moderate, and severe), depth of injury (0–3: none, mucosal, mucosal and submucosal, and transmural), and crypt damage (0–4: none, basal one-third damaged, basal two-thirds damaged, only surface epithelium intact, entire crypt, and epithelium lost). The histological score for each mouse was a sum of the score of neutrophil infiltration, depth of injury and crypt damage.

### Gut Microbiota Analysis

Fresh fecal pellets were collected in a clean sterile eppendorf tube, immediately frozen into liquid nitrogen, and then stored at −80° C. DNA was isolated using E.Z.N.A. Stool DNA Kit (Omega Bio-Tek) according to the manufacturer’s instructions. Fecal DNA samples were amplified by PCR using barcoded primer pairs targeting the V3-V4 region of 16S rRNA gene. PCR amplicons were sequenced using Illumia Mi-Seq sequencer. Bioinformatic analysis was done by Vazyme Biotech Co., Ltd., Nanjing, China. Briefly, the resulting bacterial sequence fragments were first clustered into Operational Taxonomic Units (OTUs) and aligned to microbial genes with 97% sequence similarity. Bacterial taxa summarization and rarefaction analyses of microbial diversity or compositional differences (dissimilarity value indicated by Unweighted UniFrac Distance) were then calculated and PCoA plots indicating compositional difference were generated accordingly with the Vegan package in R software.

### Accession number

The 16S rRNA sequencing data has been deposited in NCBI SRA database. The accession number is: SRP149682.

### Statistical Analysis

The data are shown as mean values ± standard error of the mean (SEM). Differences between multiple groups were compared using one-way ANOVA with post-hoc Turkey’s Multiple Comparison Test and two-way ANOVA with post-hoc Bonferroni posttests. A Student’s t-test was used for comparisons between two groups. Mantel-Cox test was used for survival analysis. Wilcoxon Signed Rank Test and Kruskal-Wallis (KW) sum-rank test were used as significance test in microbiota analysis. A P-value < 0.05 was considered significant.

## Results

### The therapeutic effect of the probiotic *B. longum JDM 301* in IBD is correlated with host microbiota

As a proof-of-concept, we used *Bifidobacterium longum* JDM 301, a widely used commercial probiotic strain in China, to test whether the therapeutic effect of probiotics is correlated to the host microbiota. For this purpose, we actively monitored the constitutes of gut microbiota in our mouse colonies by 16S rRNA sequencing. We found two different batches of C57Bl/6 wild-type (WT) mice purchased from outside resources had significant differences in their gut microbial communities (Fig. 1A). We treated them with *B. longum* JDM 301 (1×10^9^ cfu/mouse) for 3 alternate days via oral gavage, and then induced colitis by providing 3% dextran sulfate sodium (DSS) in the drinking water for 7 consecutive days and then switched back to normal water (Fig. 1B). In the absence of probiotics treatment, DSS alone induced about 15% body weight loss on day 10 in the mice from cohort A, whereas it induced around 30% body weight loss in the mice from cohort B (Fig 1C). Furthermore, without probiotics, all the mice in cohort A survived the intestinal injury-induced wasting disease; while 7 out of 8 mice in cohort B died within the experimental period. Thus, mice with different microbiota can have different sensitivity to gut epithelial injury-induced colitis. These results are consistent with the notion that the intestinal bacterial flora contributes to the immunopathogenesis of IBD (10).

**Figure 1.**
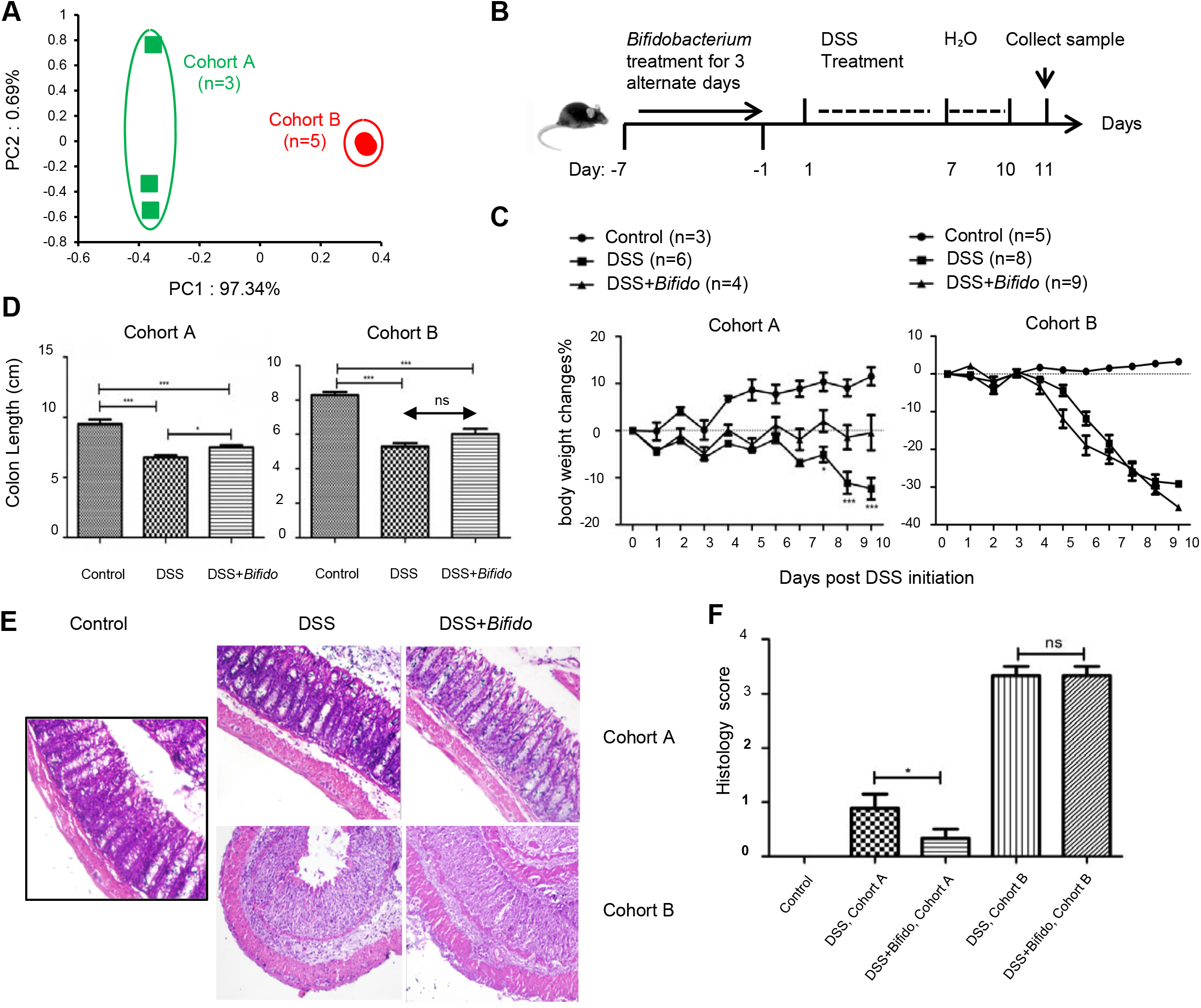
The therapeutic effect of the probiotic *B. longum* JDM 301 is correlated with host microbiota. **(A)** principal co-ordinate analysis (PCoA) based on OTU abundance of each mouse via 16S rRNA sequencing, which indicates the overall microbiota similarities between different groups. Each symbol represents one individual mouse. **(B)** Schematic diagram of experimental design. **(C)** The body weight changes during DSS treatment. **(D)** Mean colon length in cm. Colons were collected on day 11 post DSS initiation. **(E)** Representative images of H&E stained distal colon tissues from indicated mice (magnification: 200x). **(F)** Histologic scores. All data are given as means±SEMs. ns, no statistic significance; *P<0.05; ***P<0.001.

After the mice were pretreated with *B. longum* JDM 301, the body weight loss was minimized and the colon shrinking was reduced in cohort A compared to those treated only with DSS, while mice in cohort B did not show any sign of colitis improvement with probiotic treatment (Fig 1C and 1D). Microscopically, colonic epithelial damage and inflammatory cell infiltration were reduced in cohort A mice that were treated with *B. longum* JDM 301, while severe epithelial damage and inflammation remained in cohort B mice (Fig 1E and 1F). The data implied that the host microbiota not only influences IBD pathogenesis, it also influences the therapeutic effect of probiotics.

### The therapeutic effect of the probiotic *B. longum* JDM 301 in IBD is not correlated with high-fat diet

The host microbiota can be easily modified by food. Consumption of high-fat diet (HFD) is regarded as one of the risk factors of IBD and several studies demonstrated that HFD exacerbates DSS induced colitis in animals (20, 21). We were wondering whether the effect of probiotics can be modified by HFD. To test this possibility, 5 to 6-week-old C57Bl/6 male mice from cohort A and B were fed with HFD for 6 weeks before DSS treatment. 6 weeks later, we collected their fecal pellets, extracted bacterial DNA and performed high throughput 16S rRNA gene DNA sequencing. Based upon unweighted UniFrac principal coordinate analysis (PCoA), HFD more or less changed mice microbiota as expected, but differences in the bacterial flora between batch A and B still remained (Fig 2A). We then challenged the mice with DSS (Fig 2B). In consistency with the previous report, the loss of body weight after DSS challenge was more pronounced in HFD-fed versus normal chow diet (ND)-fed mice, especially in those from cohort A. Half of the mice under HFD in cohort A died after DSS challenge (Fig 2D), while none of them died under ND condition. The body weight loss and intestinal inflammation became similar between batch A and B mice (Fig 2C, 2E, and 2F), However, the mice from batch B were still more sensitive to DSS-induced wasting disease compared to mice from batch A under HFD condition (100% death rate in batch B versus 50% death rate in batch A) (Fig 2D). Overall, these data confirmed that HFD exacerbates experimental IBD as described previously (22).

**Figure 2.**
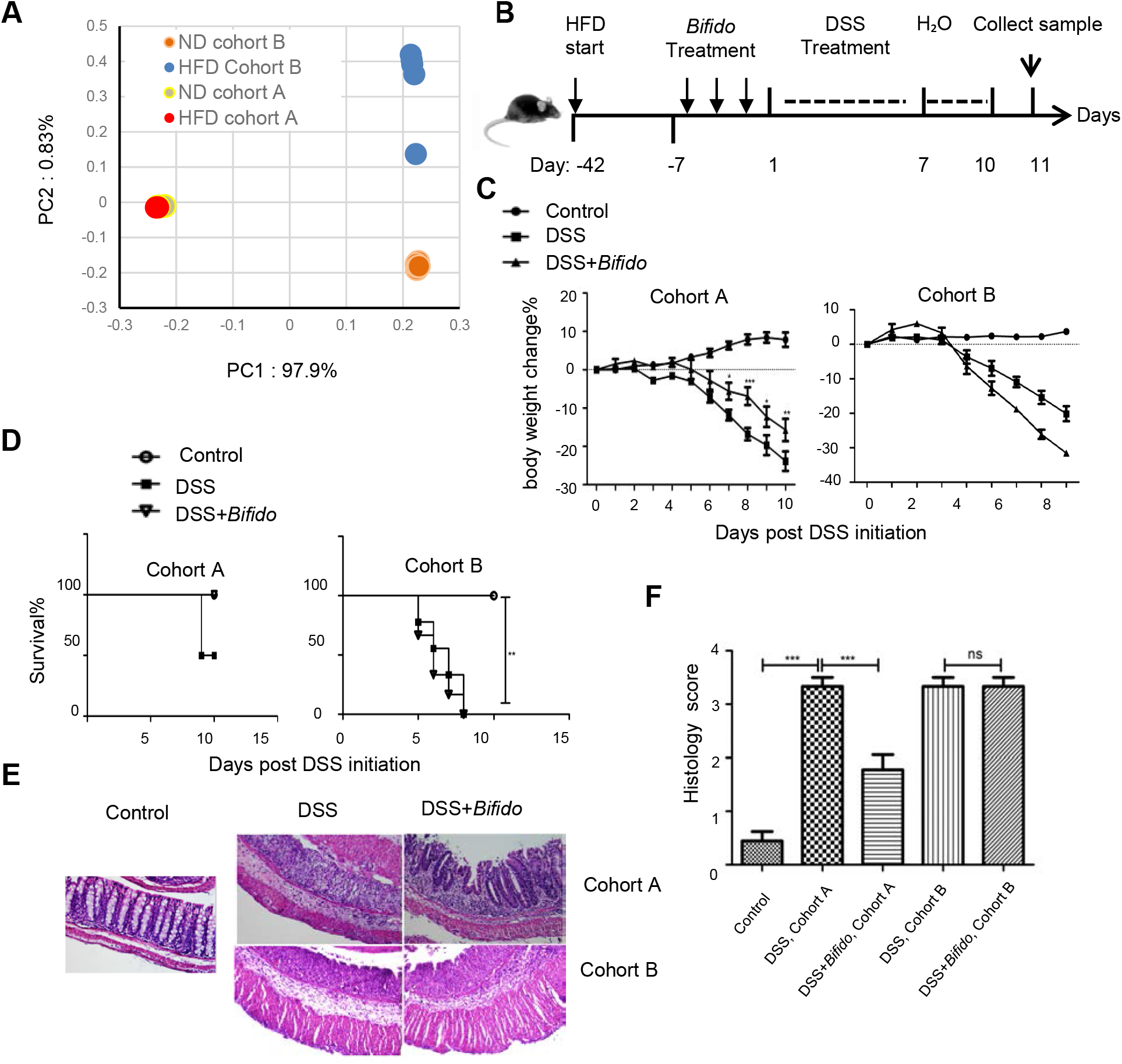
The therapeutic effect of the probiotic *B. longum* JDM 301 is not correlated with high-fat diet. **(A)** PCoA analysis illustrating the presence of different microbial community between HFD-fed mice from cohort A and B. Each symbol represents one individual mouse. **(B)** Schematic diagram for experimental design. **(C)** Body weight changes during DSS treatment in indicated mice. **(D)** Survival curve. **(E)** Representative images of H&E stained distal colon tissues from indicated mice (magnification: 200x). **(F)** Histologic scores. All data are given as means±SEMs. *P<0.05; **P<0.01; ***P<0.001. The number of mice per group was 3~6.

To examine whether changes in diet can result in alteration of the therapeutic effects of the probiotics, the different batches of mice that fed on HFD were orally gavaged with *B. longum* JDM 301 one week before DSS challenge. Similar to the results obtained under the ND condition, the therapeutic effect of *B. longum* JDM 301 still was different between cohort A and B in HFD-fed condition. The body weight loss after DSS challenge was significantly relieved in mice from cohort A but not cohort B when treated with probiotics (Fig 2C). The colonic inflammation also became less severe in cohort A but not cohort B (Fig 2E and 2F). The result indicated that although HFD can exacerbate colitis, it may not be able to determine the outcomes of probiotic therapeutic effect in IBD. It is likely that certain preformed postnatal microbial members that are not disturbed by high-fat diet control the probiotic effectiveness.

### Ecological characteristics of the gut microbiota that are correlated with *B. longum* JDM 301 efficacy

To look for the ecological characteristics of the gut microbiota that are correlated with the probiotic’s effectiveness, we compared the overall community configurations in probiotic-sensitive cohort A and probiotic-insensitive cohort B mice at both ND and HFD-fed conditions. Significant differences in both species richness represented by Chao1 index and species evenness represented by Shannon’s index were observed between cohort A and cohort B mice (Fig 3A). Both indices were bigger in cohort B mice compared to those in cohort A mice, irrespective of the diet used. HFD feeding reduced total species richness (Chao1) in both cohorts, but species evenness (Shannon index) was not disturbed by HFD (Fig 3A). At phyla level, cohort A mice had more Firmicutes, Actinobacteria, Saccharibacteria and less Bacteroidetes compared to cohort B at both ND or HFD conditions (Fig 3B). Shifting diet from ND to HFD resulted in increase of the phylum Proteobacteria in both cohorts (from 1 and 5% to 14%, respectively) (Fig 3B). This is consistent with the notion that Proteobacteria expansion is an indicator of colon epithelial dysfunction and correlates to the increase of sensitivity to DSS-induced colitis at HFD condition (23). However, Proteobacteria was not a good indicator for the *B. longum* JDM 301’s efficacy, as the abundance of Proteobacteria were not different between cohort A and B at both ND or HFD condition (Fig 3B). Furthermore, 18 different genera were found to be consistently different between the two cohorts of mice irrespective of the diet used (Fig 3C). Among them, *Alistipes* and *Parabacteroides*, two genera that have been implied participating in IBD pathogenesis (24), were increased significantly in mice from cohort B fed with either ND or HFD.

**Figure 3.**
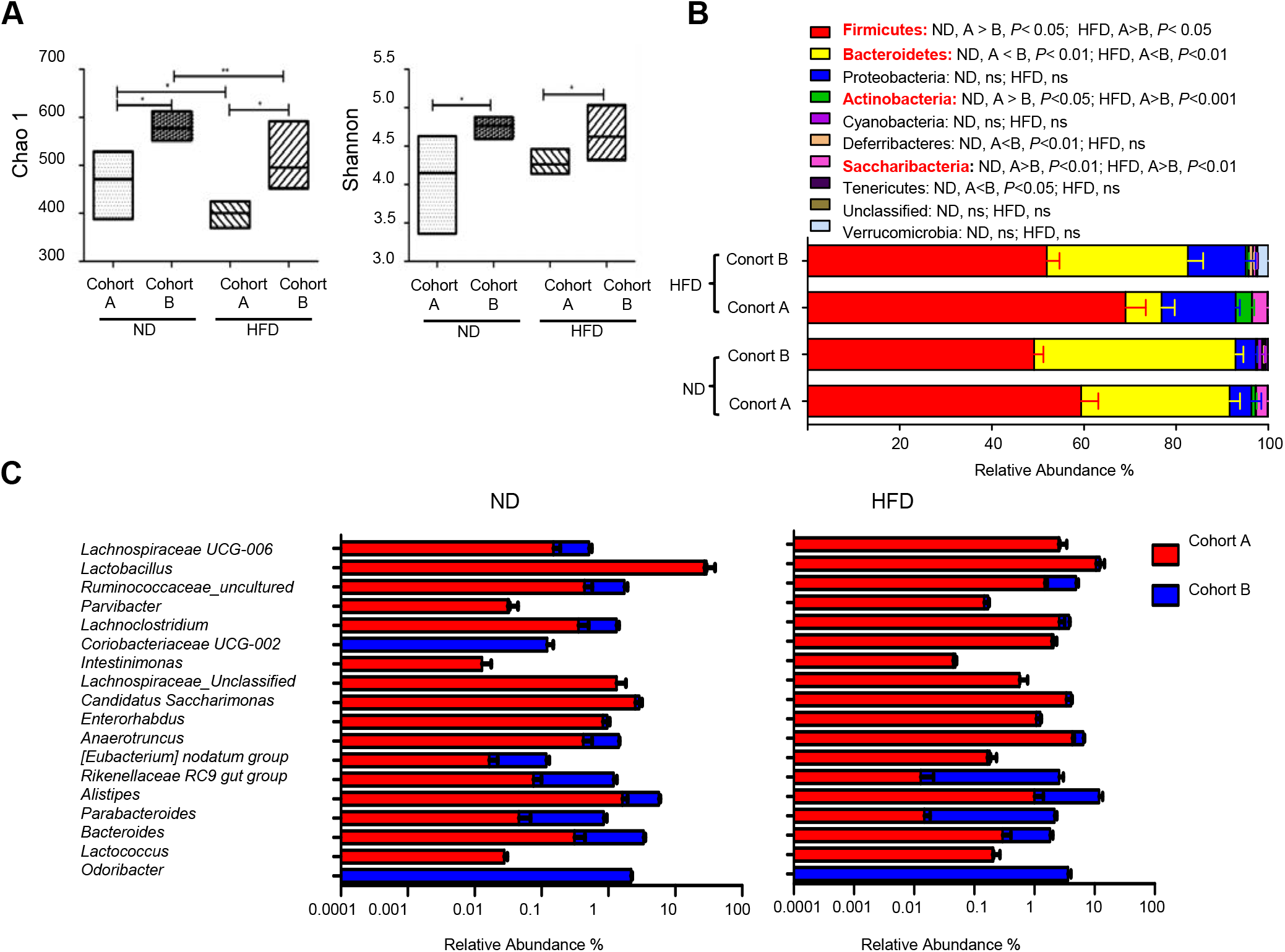
Ecological characteristics of the gut microbiota that are correlated with *B. longum* JDM 301 efficacy. **(A)** α-diversity indicated by Chao1 (species richness) and Shannon index (species evenness). The line drawn in the middle of the box represents the median value and the box represents the range of values. **(B)** Taxonomic composition at the phyla level in the indicated mice. **(C)** Taxonomic composition at genera level in the indicated mice under ND and HFD. The top 18 genera that were significantly different (P<0.05) between the two cohorts in both ND and HFD conditions were shown. *P<0.05; **P<0.01.

### *B. longum* JDM 301 has limited ability to change the taxonomic composition of the gut microbiota

The mechanism of how probiotics work remain largely unknown. One possibility is that the probiotics change the host bacterial flora. To determine if probiotic *B. longum* JDM 301 alters the microbiome, we performed high-throughput gene-sequencing analysis of 16S rRNA in fecal bacterial DNA isolated from probiotic untreated and treated WT mice from batch A one day before DSS challenge. We used rarefaction analysis to compare bacterial diversity within individual mice of a group (α diversity) in both ND- and HFD-fed conditions. *B. longum* JDM 301 treatment did not change species richness (Chao1) (Fig 4A and 4B) and species evenness (Shannon index) significantly (Fig 4C and 4D). PCoA analysis of the microbiota composition in probiotic treated mice did not show a different community composition relative to that of probiotic untreated mice in both ND- and HFD-fed conditions (Fig 4E). Thus, the impact of *B. longum* JDM 301on the taxonomic composition of the fecal microbiota was very limited.

**Figure 4.**
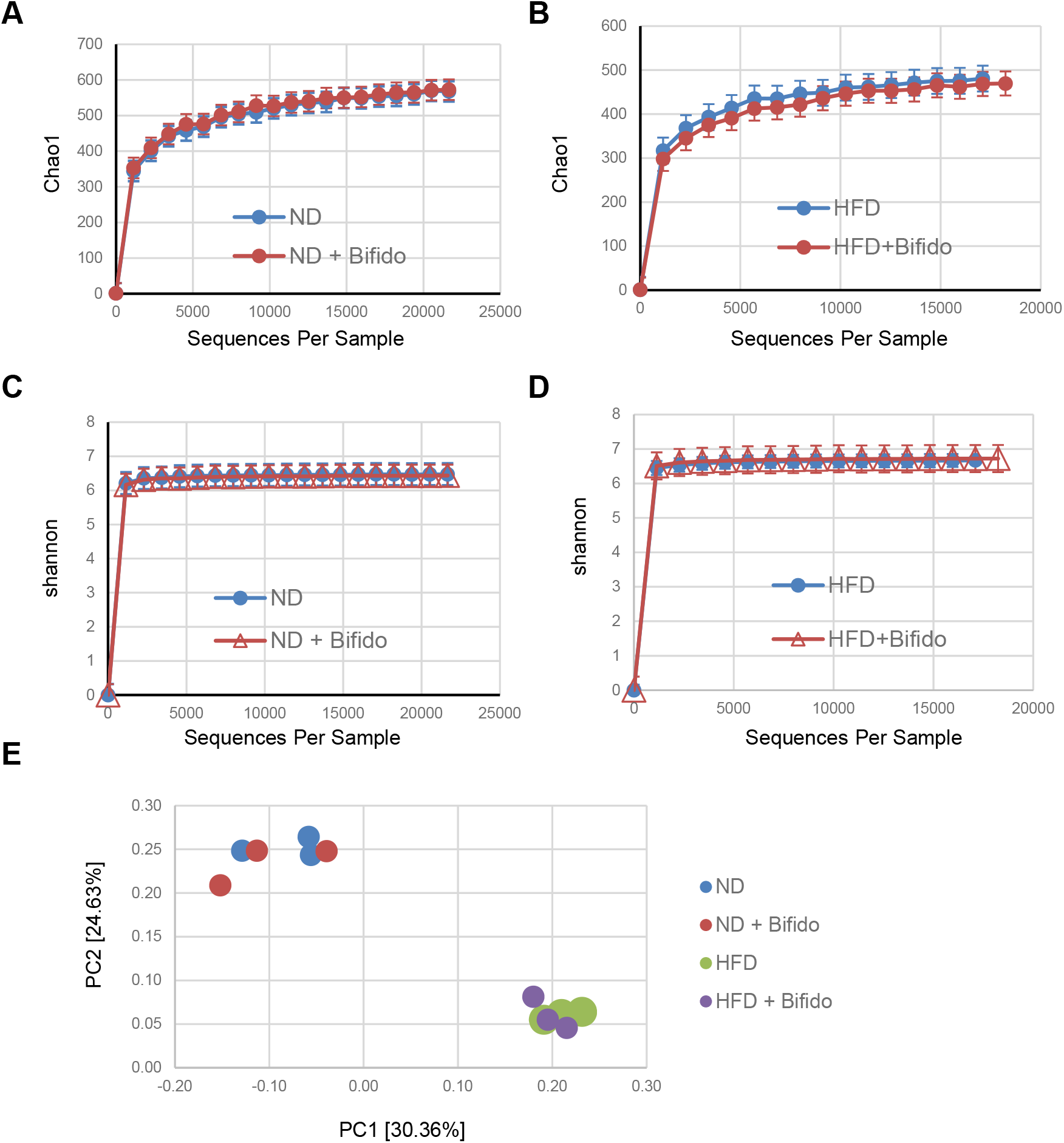
*B. longum* JDM 301 has limited ability to change the taxonomic composition of the gut microbiota. **(A to D)** High-throughput sequencing of 16S rRNA in fecal bacterial DNA from WT mice in batch A fed with ND or HFD. Chao1, indicative of bacterial species richness (A and B), Shannon, indicative of bacterial species evenness (C and D). **(E)** PCoA analysis of the microbiota composition in indicated mice. Each symbol represents one individual mouse.

### The therapeutic effect of the probiotic *B. longum* JDM 301 in IBD requires TLR2 signals

Another possibility of how probiotics work in IBD pathogenesis is to engage the host cells, particularly the host immune system to maintain intestinal homeostasis. Toll-like receptors (TLRs) are critical host sensors for microbes. In a rat necrotizing enterocolitis model, the effect of *Bifidobacterium bifidum* to reduce mucosal injury and to preserve intestinal layer was reported to be through the TLR2 pathway (25). We posited that the effectiveness of the probiotic *B. longum* JDM 301 in treating IBD might also depend on TLR2 signals. To test this hypothesis, colitis was induced in *TLR2^-/-^* mice fed with either ND or HFD. In consistency with previous report that TLR2 plays critical role in maintaining gut epithelial homeostasis (26, 27), *TLR2^-/-^* mice raised in our facility also developed severe DSS-induced colitis indicated by severe body weight loss, high histologic scores, and 100% mortality rate in both ND- and HFD-fed conditions (Fig 5A-5D). To determine if probiotic therapy could ameliorate DSS-induced colitis in *TLR2^-/-^* mice, the mice were pretreated with *B. longum* JDM 301 and challenged with 3% DSS. Unlike the WT mice in cohort A, *TLR2^-/-^* mice pretreated with *B. longum* JDM 301 did not show improvement in body weight loss (Fig 5A). No difference was obtained on the survival rate with or without *B. longum* JDM 301 treatment (Fig 5B). All the *TLR2^-/-^* mice challenged by DSS died before the end of the experiment. Severe colonic inflammation evaluated by H&E stained samples remained the same with or without *B. longum* JDM 301 treatment (Fig 5C and 5D). The data imply a requirement of intact TLR2 signals in establishing the protective effect by *B. longum* JDM 301.

**Figure 5.**
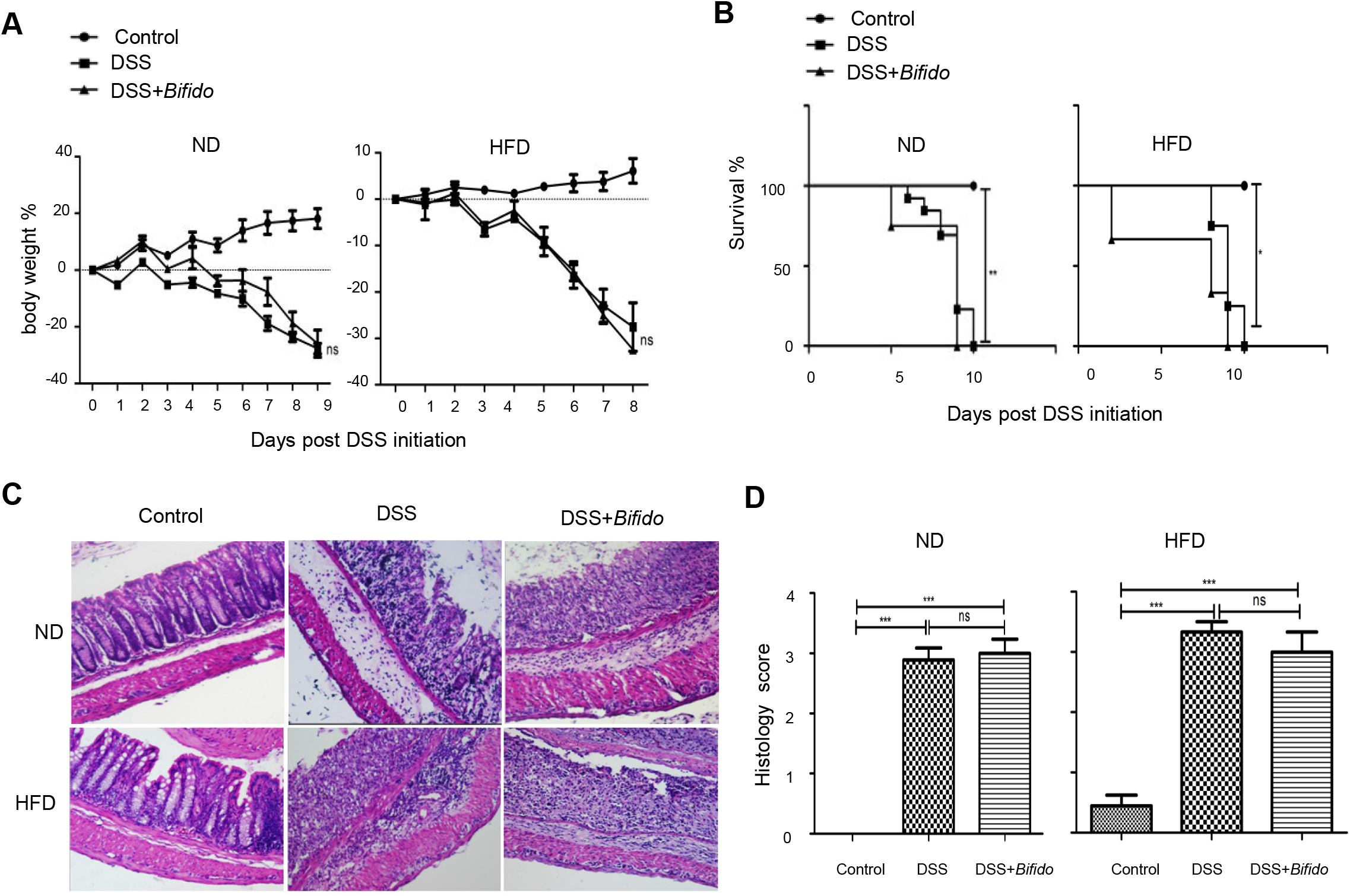
The therapeutic effect of the probiotic *B. longum* JDM 301 in IBD requires TLR2 signals. **(A)** Body weight changes during DSS treatment in *TLR2:^-/-^* mice that feed on ND or HFD. **(B)** Survival curve. **(C)** Representative images of H&E stained distal colon tissues from indicated mice (magnification: 200x. **(D)** Histologic scores. All data are given as means±SEMs. *P<0.05; **P<0.01; ***P<0.001. The number of mice per group was 3~6.

## Discussion

In this work, we measured the ability of gut microbiota, high-fat diet and host genetic factor (e.g. TLR2) to influence the host response to a model probiotic, *B. longum* JDM 301, in a DSS-induced mouse colitis model. We demonstrated that the probiotic therapeutic effect can be varied across individual mouse even when the mice have the same genetic background and consume the same type of food. We further showed different microbiome features were highly correlated with different probiotic response. Consumption of diet rich in fat can change the host sensitivity to mucosal injury-induced colitis, but may not necessarily change the host responsiveness to probiotic therapy. Finally, the host genetic factor TLR2 was also required for a therapeutic effect of *B. longum* JDM 301.

Although probiotics are defined as beneficial microorganisms to the host, exact mechanisms of how probiotics function between the host and the gut microbiome remain incompletely understood. The bacterial species that can be called probiotics are still expanding (28), but whether one type of probiotic fits for all people at the same or different disease conditions is currently not clear. Our data suggested that the individual host gut microbiome can probably influence whether a given probiotic can have beneficial effects on the specific host or not. Possible pathways that have been suggested for how probiotics works include: (i) restoring microbial imbalances, (ii) enhancing the epithelial barrier function and/or (iii) modulating the immune responses (29). It remains to be determined which pathways can be modified by the host microbiome.

How personalized microbiome influence probiotic effect requires further investigation. One possible influence of the host microbiome is to influence the probiotic engraftment efficacy (30–32). Another possible influence of the microbiome to the probiotics is to influence their functions. One earlier study indicated that when the gut microbes translocated to the internal tissue, they can induce disease tolerance (33). Many other more possible mechanisms remain to be determined.

Our data further suggested that it might be possible to predict probiotic efficacy via analysis of the host microbial and genetic features. Personalized measurements including gut microbiome have been shown to be able to more accurately predict postprandial glycemic response for each unique person (34), it might also help for personalized probiotic therapies.

In aggregate, this study demonstrated correlation of individual host microbiome and genetics to the protective effects of probiotic therapy in colitis. Therefore, carefully monitoring personal and microbiome features might be needed for a success of probiotic therapy for IBD patients.

## ACKNOWLEDGEMENTS

Project support was provided in part by the National Natural Science Foundation of China (Grant Nos. 81770853 to Y.W.), the Priority Academic Program Development of Jiangsu Higher Education Institutions (PAPD) in the year of 2014 (Grant No. KYLX14-1448), Natural Science Foundation of the Higher Education Institutions of Jiangsu Province, China (Grant Nos 16KJB310016), the Starting Foundation for Talents of Xuzhou Medical University (Grant No. D2016029 to Y.W.).

## Author Contributions

Conceived and designed the experiments: Y. Wang, K. Zheng; Performed the experiments: S. Suwal, Q. Wu, W. Liu, Q. Liu, H. Sun, M. Liang, J. Gao, Y. Kou, Z. Liu, Y. Wei; Analyzed the data: Y. Wang, S. Suwal; Wrote the paper: Y. Wang, S. Suwal.

